# Stimulus selection drives value-modulated somatosensory processing in superior colliculus

**DOI:** 10.1101/2024.06.28.601226

**Authors:** Yun Wen Chu, Suma Chinta, Hayagreev V.S. Keri, Shreya Beri, Scott R. Pluta

## Abstract

A fundamental trait of intelligent behavior is the ability to respond selectively to stimuli with higher value. Where along the somatosensory hierarchy does information transition from a map of stimulus location to a map of stimulus value? To address this question, we recorded single-unit activity from populations of neurons in somatosensory cortex (S1) and midbrain superior colliculus (SC) in mice conditioned to respond to a positive-valued whisker stimulus and withhold responses using an adjacent, negative-valued whisker stimulus. The stimulus preference of the S1 population was equally weighted towards either whisker, in line with a somatotopic map. Surprisingly, we discovered a large population of SC neurons that were disproportionately biased towards the positive stimulus. This disproportionate bias was controlled by spike facilitation for the positive stimulus and spike suppression for the negative stimulus in single neurons. Removing the opportunity for mice to select the positive stimulus reduced stimulus bias in SC but not S1, suggesting that sensory processing in SC neurons was partially controlled by movement preparation. Similarly, the spontaneous firing rates of SC but not S1 neurons accurately predicted reaction times, suggesting that SC neurons play a persistent role in perceptual decision-making. Taken together, these data indicate that the somatotopic map in S1 is transformed into a value-based map in SC that encodes stimulus priority.

## Introduction

Imagine that it is raining outside, and gusts of wind are ripping through the trees. While looking out the window, you approach the closet to find a jacket. You quickly push aside a cotton t-shirt before grabbing a parka and heading out the door. In daily life, we encounter countless objects in our environment. To optimize survival, we select the objects that add value to our life and dismiss the objects that add unnecessary costs. Guiding this behavior are maps of physical space in our brain that are repeated across the different stages of sensory processing. Where along this hierarchy does information transition from a map faithful to physical location to a map interwoven with object value? To address this question, we utilized the somatosensory whisker system of mice, where a map of stimulus space is clearly organized across the different stages of sensory processing [1–8].

In this study, we focused on the primary sensory cortex (S1) and midbrain superior colliculus (SC), due to their known importance in sensory guided behaviors [9–12]. Both brain areas receive bottom-up whisker information that encodes the location of stimuli in somatotopic coordinates. Both areas receive top-down input from motor and association cortices, providing a circuit basis for contextualizing stimulus information with goal-oriented actions [13–16]. S1 provides monosynaptic sensory drive to SC, and ascending SC projections to the thalamus augment whisker responses in S1, forming a reciprocal flow of tactile information [13,17,18]. Despite these known relationships, how sensory associations differentially shape their representations of space is unclear. While each brain area has been studied in isolation, there is no study comparing somatosensory processing in each area during the same goal-oriented task.

Behavioral context is known to modify neuronal receptive fields in S1 and SC. For example, while performing a shape discrimination task that involves multiple whiskers, S1 neurons located in the most task-relevant barrel column expand their receptive fields [19]. In addition, S1 neurons adapt their tactile sensitivity to match the intensity of the task-relevant stimulus distribution [20]. In SC, neuronal receptive fields expand and increase their sensitivity when the animal is prepared to make a saccadic eye movement [21,22], suggesting that movement preparation enhances SC visual processing. Therefore, receptive field properties in S1 may adapt to optimally encode the task-relevant stimulus features, while sensory processing in SC appears to have a stronger relationship to orienting movements and decision-making.

To determine how the representation of sensory space in S1 and SC is modulated by behavioral context, we recorded single-unit activity from populations of neurons in mice performing stimulus selection guided by value-based associations. In one version of the task, two adjacent single-whisker stimuli had opposite values (positive/negative) and elicited opposing behavioral responses (Go/No-Go). In the other version, the two single-whisker stimuli had equal value (positive) and elicited the same behavioral response (Go). In both versions of the task, we found that the stimulus preference of the S1 population was equally weighted towards either whisker stimulus. Therefore, stimulus value and its associated action did not modify the representation of sensory space in S1. Conversely, we found that behavioral context strongly influenced the representation of whisker space in SC. SC neurons were strongly biased towards the positive valued stimulus, but only when the stimuli and their associated actions had opposite values. When the stimuli had equal value, stimulus bias in the SC population became smaller than S1. Removing the opportunity for mice to select the positive valued stimulus reduced bias in SC, but not S1. Likewise, the spontaneous firing rate of SC but not S1 neurons predicted the speed of perceptual decision-making. Taken together, these data suggest that S1 contains a faithful somatotopic map of whisker space that is transformed into a value-based and action-sensitive map in SC that encodes stimulus priority.

## Results

To determine the impact of stimulus value on somatosensory processing in S1 and SC, we trained mice to discriminate between two adjacent single-whisker stimuli of opposing values. Mice performed the task by touching an object that entered the movement field of either the positive-valued “Go” whisker or the negative-valued “NoGo” whisker (Fig. 1A, B). Mice reported the presence of the positive-valued stimulus by licking a port to receive water reward (Hit response, Suppl. Fig. 1). If mice licked the water port during presentation of the negative-valued stimulus (False alarm), they were forced to locomote 2 – 4 times the normal distance to start the next trial. Mice touched the positive and negative objects with equal frequency, and in nearly all trials (92%) responded with a lick latency longer than 0.3 seconds from first touch (Fig. 1C). While mice performed the task, we tracked whisker movement and recorded single-unit activity from populations of S1 or SC neurons using a high-density, three-shank silicon probe (Fig. 1D). We targeted our electrodes to the SC whisker map using previously established stereotaxic coordinates and validated our placement by manually deflecting individual whiskers (Fig. 1E) [17,23]. S1 recordings were guided to the whisker columns involved in the task using intrinsic signal imaging, with somatotopic locations confirmed post-hoc by examining touch-triggered local field potentials across the electrode shanks (Fig. 1F & Suppl. Fig. 1E). Aligning S1 and SC spiking to the onset of the first touch in each trial revealed short latency tactile responses with varying rates of adaptation and a preference for either the positive or negative whisker (Fig. 1G, H & Suppl. Fig. 1F). A total of 11 mice were trained to perform this version of the task. Most SC (76%, 549/727) and S1 (89%, 625/705) neurons displayed a significant tactile response to either or both stimuli (p < 0.05, 1-way ANOVA with Tukey comparison).

**Figure 1.**
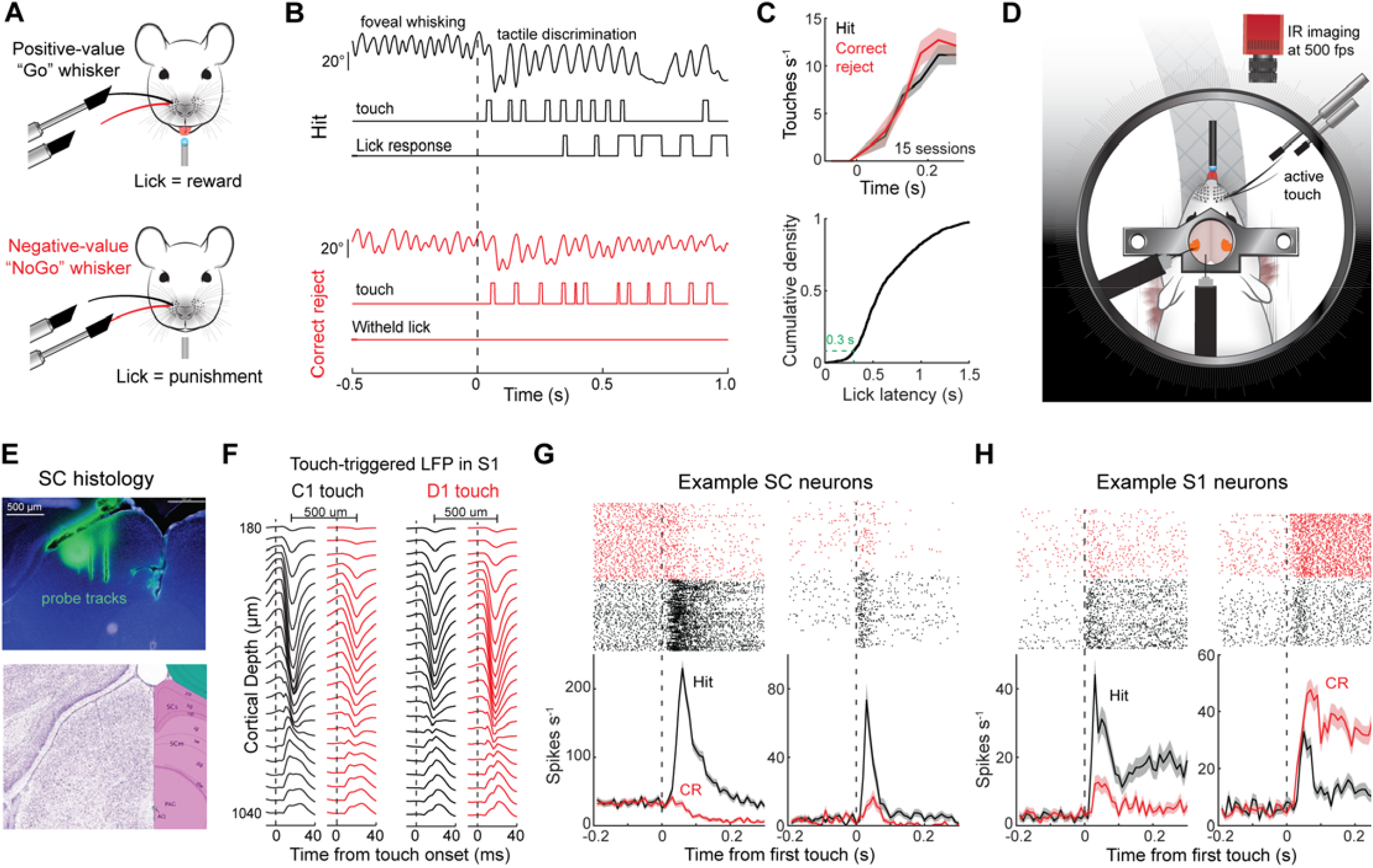
The formation of value-based sensory associations using the somatosensory whisker system. **A**) Illustration of the positive- and negative-valued single whisker stimuli. A lick response to the positive stimulus was associated with reward while a lick response to the negative stimulus was associated with punishment. **B**) Whisker angle, touch duration, and lick responses during example Hit and Correct Rejection (CR) trials. **C**) Top, mean touch rate during the Hit and CR trials across 15 behavioral sessions in 11 mice. Bottom, cumulative density of lick latencies (1655 trials across 15 sessions in 11 mice). **D**) Illustration of the experimental system combining active tactile discrimination, whisker tracking, and electrophysiology in either the barrel cortex or superior colliculus. **E**) Top, histological section of superior colliculus and three-shank silicon probe dye labeling. Bottom, Allen Brain Atlas of corresponding region of superior colliculus. **F**) Touch-triggered local field potential (LFP) across two electrode shanks in somatosensory cortex (S1) during C1or D1 touch. Notice the change in LFP amplitude between the two stimuli, indicating a somatotopic map. **G**) Rasters and histograms of firing rate in two SC neurons on trials when the animal responded to the positive stimulus (Hit, black), and when it withheld a response to the negative stimulus (CR, red). **H**) Same as in G, except for two example S1 neurons.

### Task structure enhances positive stimulus bias in SC neurons

Both SC and S1 contain a somatotopic map of whisker space, where each whisker on the face activates a corresponding coordinate in each brain area [4,5,7]. We reasoned that if a brain area is faithful to this map, then the stimulus preference of its neural population would be evenly divided among both whiskers. In line with this reasoning, we found that the stimulus preference of the S1 population was evenly distributed, reflecting only a small sampling bias (58%) towards the negative whisker (Correct Rejection (CR), Fig. 2A). However, the stimulus preference of the recorded SC population was remarkably skewed (70%) towards the positive whisker (Hit). To determine if the strength of stimulus bias in SC and S1 neurons changed according to their stimulus preference (positive vs. negative whisker), we compared the Hit and CR responses within each group (Fig. 2B, D). Among neurons that preferred the positive stimulus, the bias of the SC population was significantly greater than the bias of the S1 population (Fig. 2C; p = 3.8e-9, 1-way ANOVA with Tukey comparison). However, among neurons that preferred the negative stimulus, bias among S1 and SC neurons was equivalent and indistinguishable from the positive preferring S1 population (Fig. 2D, E, p > 0.9, 1-way ANOVA with Tukey comparison). Therefore, SC neurons had a uniquely large positive stimulus bias.

**Figure 2.**
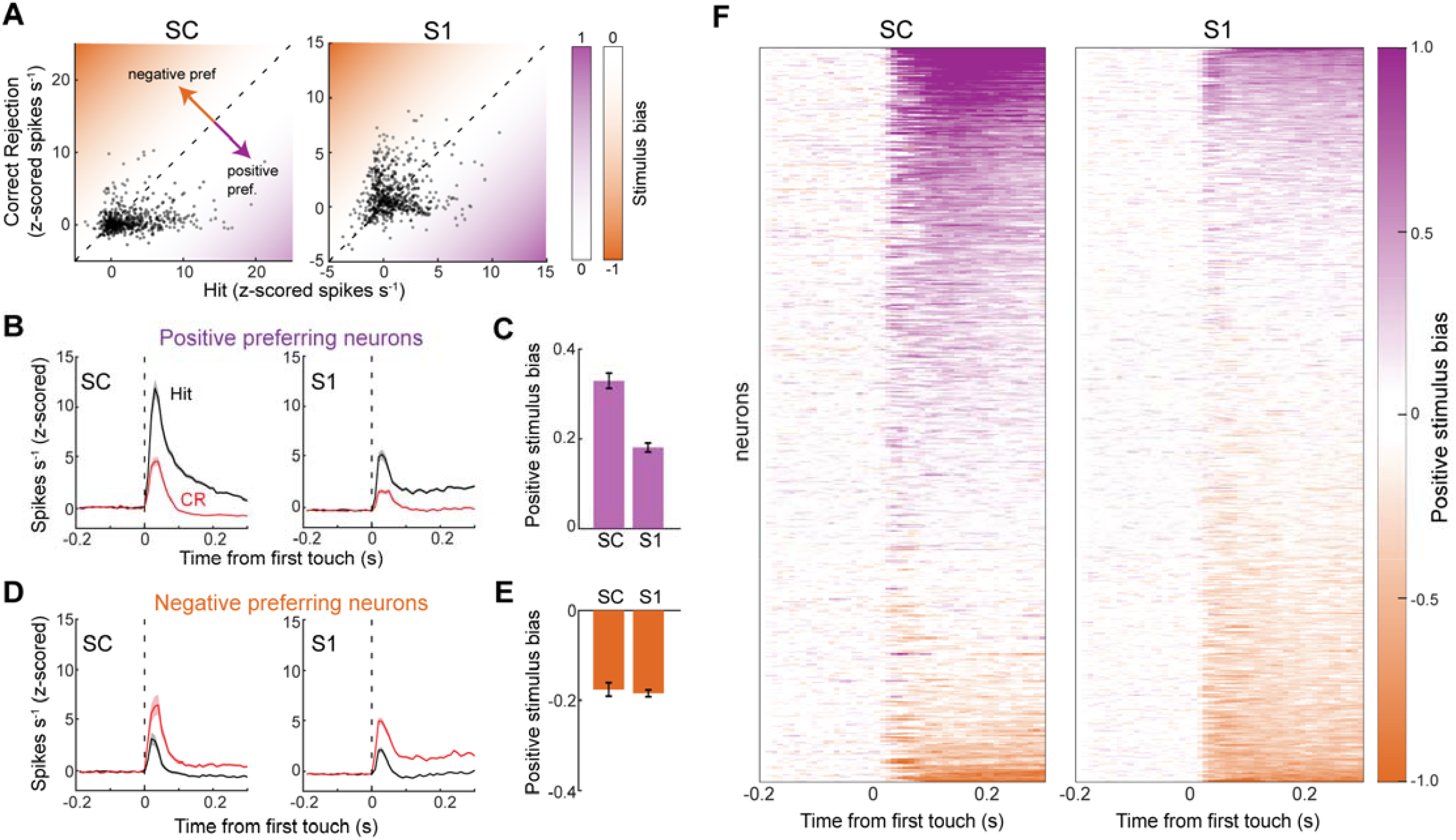
SC neurons are strongly biased towards the positive stimulus. **A**) Scatter plot comparing neuronal firing rates (z-scored) during Hit and Correct Rejection (CR) trials (549 neurons in 8 mice in SC; 625 neurons from 7 mice in S1). **B**) Histograms of firing rate during Hit and CR trials in neurons that preferred the positive stimulus (386 neurons in 8 mice in SC; 263 neurons from 7 mice in S1). **C**) Mean positive stimulus bias in positive preferring SC and S1 neurons (p = 7.7e-11, rank sum test). **D**) Histograms of firing rate (z-scored) during Hit and CR trials in neurons that preferred the negative stimulus (163 neurons in 8 mice in SC; 362 neurons from 7 mice in S1). **E**) Mean positive stimulus bias in negative preferring SC and S1 neurons (p = 0.01, rank sum test). **F**) Heat maps of positive stimulus bias across the SC and S1 populations.

Next, we hypothesized that positive bias in SC was created by the opposing stimulus values underlying the task structure. To test this possibility, we trained a separate cohort of mice to detect and associate both whisker stimuli with reward, causing the two stimuli to have equal value and elicit equivalent behavioral responses (Fig. 3A & Suppl. Fig. 2). A total of 6 mice were trained for this task, of which six SC and four S1 recording sessions were performed. The majority of SC (67%, 332/449) and S1 (76%, 353/466) neurons displayed a significant tactile response to either or both stimuli (p < 0.05, 1-way ANOVA with Tukey comparison). Interestingly, we now found that the stimulus preference of the SC and S1 populations was evenly distributed between both whiskers (Fig. 3B; SC: 52% whisker 1 preferring; S1: 42% whisker 1 preferring). Moreover, stimulus bias of the SC population was now slightly smaller than S1 (Fig. 3C – G, p = 0.005, rank sum test, bias for both whisker stimuli combined). Taken together, these data reveal that stimulus bias in SC, but not S1, was enhanced by the process of preferentially selecting one stimulus while ignoring the other.

**Figure 3.**
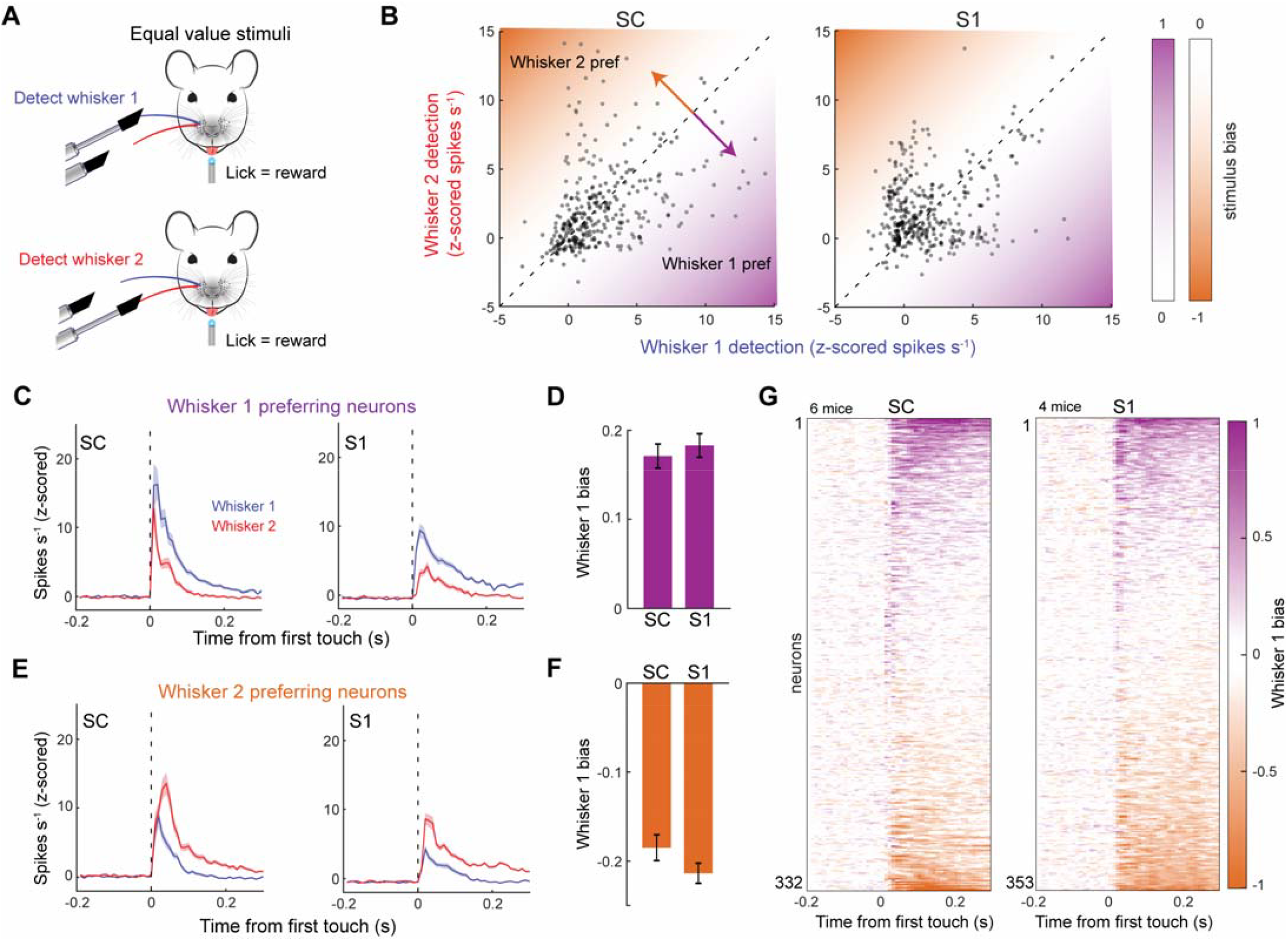
Equal valued sensory associations reduce bias in SC neurons. **A**) Illustration of the task structure. Licking in response to either of the single-whisker stimuli was paired with reward. **B**) Scatter plots comparing neuronal firing rates (z-scored) during the detection of the two different single whisker stimuli (332 neurons in 6 mice in SC; 353 neurons from 4 mice in S1). **C**) Histograms of firing rate (z-scored) in neurons that preferred the whisker 1 stimulus (172 neurons from 6 mice in SC; 149 neurons in 4 mice in S1). **D**) Whisker 1 stimulus bias in neurons that preferred the whisker 1 stimulus. **E**) Histograms of firing rate (z-scored) in neurons that preferred the whisker 2 stimulus (160 neurons in 6 mice in SC; 204 neurons in 4 mice in S1). **F**) Whisker 1 stimulus bias in neurons that preferred the whisker 2 stimulus. **G**) Heat maps of whisker 1 bias among the population of S1 and SC neurons.

### Spike facilitation and suppression enhance stimulus bias in SC neurons

Next, we sought to identify the neural dynamics underlying the large positive stimulus bias in SC neurons. First, we noticed that the CR response in SC neurons rapidly changed over time, with firing rates initially increasing but then quickly decreasing, often to a level below the pre-stimulus baseline (Fig. 4A). This delayed suppression of the CR response coincided with an increasing stimulus bias in SC; however, in S1, stimulus bias was constant over time (Fig. 4B, C). To clearly delineate the relationship between neural dynamics and stimulus bias, we quantified the bias of the Hit and CR responses relative to the pre-stimulus baseline. In the SC population, the CR response was initially positive but then decreased at a rate much faster than the Hit response (Fig 4D). This rapid decrease in the CR response became negative ∼70 ms post first touch and reached its asymptote ∼50 ms later, which coincided with the largest positive stimulus bias in the SC population (shown in Fig. 4C). Conversely, in the S1 population, the Hit and CR responses increased and decreased at similar rates, leading to a constant stimulus bias over time (shown in Fig. 4C). This indicates that sustained spike facilitation during Hit trials combined with spike suppression during CR trials enhances positive stimulus bias in SC neurons. Stimulus bias over time and the underlying neural dynamics of negative preferring neurons was similar between SC and S1 (Suppl. Fig. 3).

**Figure 4.**
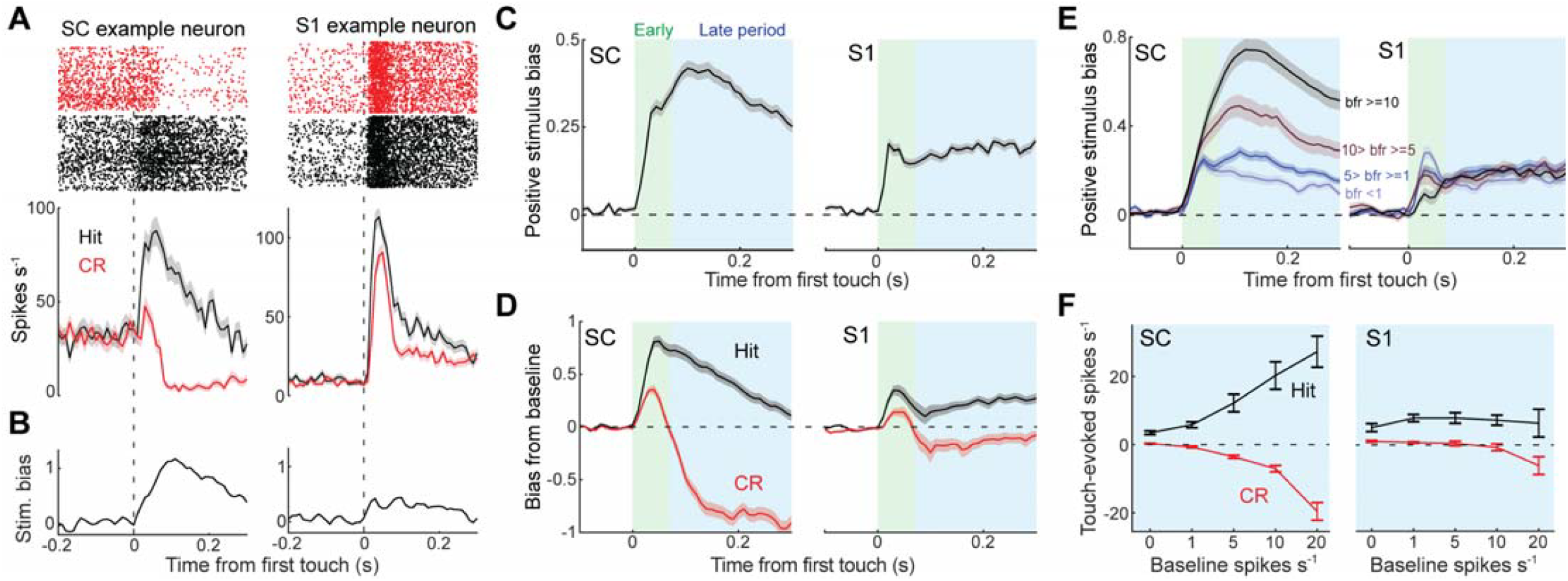
Stimulus bias in SC is controlled by sustained spike facilitation and delayed spike suppression in single neurons. **A**) Rasters and histograms of mean firing rate in an example SC and S1 neuron during Hit and Correct Rejection (CR) trials. **B**) Positive stimulus bias in the same example neurons. **C**) Mean positive stimulus bias relative to first touch among the positive preferring SC and S1 neurons (386 neurons in 8 mice in SC; 263 neurons from 7 mice in S1). **D**) Mean bias relative to baseline firing rate during Hit and CR trials in positive preferring SC and S1 neurons (190 neurons in 8 mice in SC; 118 neurons from 7 mice in S1; Showing neurons with baseline spike rate > 4). **E**) Mean positive stimulus bias in positive preferring SC and S1 neurons where the populations are sorted by baseline firing rate (For SC: 107 neurons in 8 mice in bfr>=10 group; 60 neurons in 8 mice in 10>bfr>=5 group; 131 neurons in 8 mice in 5>bfr>=1 group; 88 neurons in 8 mice in bfr<1 group; For S1: 60 neurons in 7 mice in bfr>=10 group; 42 neurons in 6 mice in 10>bfr>=5 group; 99 neurons in 7 mice in 5>bfr>=1 group; 62 neurons in 5 mice in bfr<1 group). **F**) Mean touch-evoked spike rates during the late period of the sensory response among populations grouped according to their baseline firing rate.

We reasoned that if spike suppression during CR trials enhances stimulus bias in SC, then neurons with a higher baseline firing rate should have a higher stimulus bias, due to their greater distance from the floor. In support of this logic, we found that baseline firing rate was directly related to positive stimulus bias in SC but not S1 neurons, particularly for the later period of the response, when spike suppression was greatest (Fig. 4E). To determine the contribution of the Hit and CR responses to this trend, we plotted their touch-evoked firing rates during the late sensory period (70 – 300 ms from first touch, Fig 4F). In the SC population, neurons with higher baseline firing rates displayed larger increases in their Hit response, while also displaying larger decreases in their CR response. Therefore, the gap between the Hit and CR responses widened in neurons with larger baseline firing rates. Conversely, in the S1 population, the gap between the Hit and CR responses did not change as a function of baseline firing rate. Taken together, these data reveal that SC neurons with a higher baseline firing rate are more strongly biased towards stimuli with a positive association between action (licking) and outcome (water reward). Is sensory processing altered when mice cannot perform the action associated with the positive outcome?

### Task engagement and movement preparation enhance SC sensory processing

The SC is long known to play a critical role in sensory-guided movements [24,25]. Given this knowledge, we hypothesized that tactile responses in SC neurons are influenced by whether or not the animal has the opportunity to select the positive stimulus. To test this hypothesis, we compared sensory responses on Hit trials to when the animal Missed (failed to respond) or was experimentally denied the opportunity (Away) to respond (Fig. 5A & Suppl. Fig. 4A – C). Miss trials were sporadic and primarily occurred in the latter half of the session, while Away (water port removed) trials were experimentally controlled and occurred at regularly spaced intervals (Fig. 5B). We focused our analysis on neurons that preferred the positive stimulus, given their strong stimulus bias and potential for influencing the decision to lick. In many SC neurons, the response to the positive stimulus was often larger on Hit than Miss (48% 133/276) and Away (38% 102/268) trials, as shown in an example neuron and in population statistics, suggesting that task engagement and movement preparation increase tactile sensitivity (Fig 5C – G). In S1, the change in sensory responsiveness on Miss trials (55% 75/136) was apparent but less consistent than SC, while the Away condition (17% 30/178) had no clear effect (Fig. 5C – G). Positive stimulus bias in SC but not S1 neurons significantly decreased on Away trials, suggesting that sensorimotor processing in SC neurons is augmented by movement preparation (Fig. 5H). Differences in touch strength and locomotion speed could not explain these effects (Suppl. Fig. 4D, E). SC neurons that were modulated by Miss were similarly modulated by Away, indicating that task engagement and movement preparation were represented by the same population of SC neurons (Fig. 5I). Sensory processing in negative preferring neurons was largely unaffected by behavioral context (Suppl. Fig. 4F – K).

**Figure 5.**
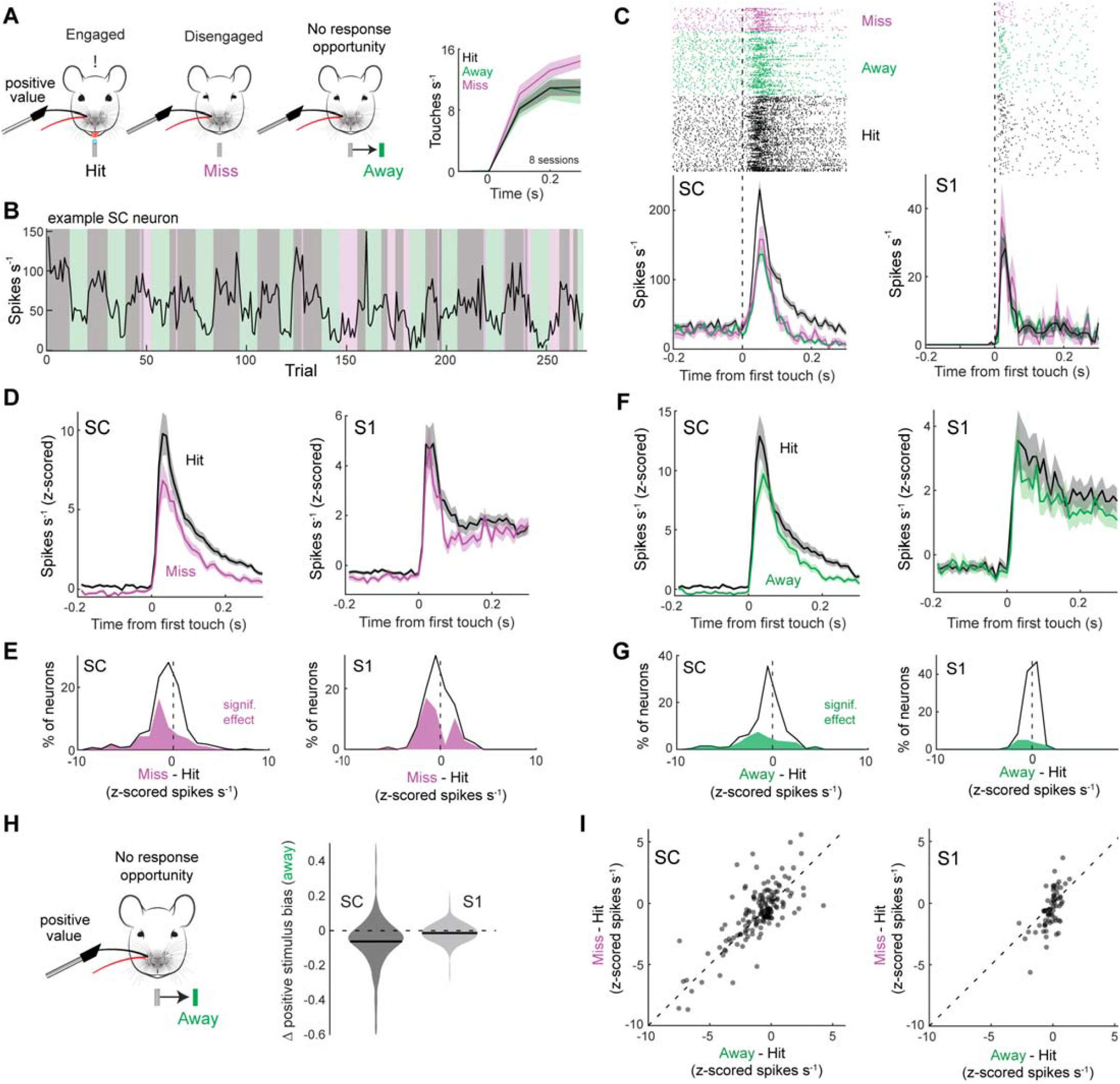
Movement preparation increases positive stimulus bias in SC neurons. **A**) Diagram illustrating the three different behavioral contexts during positive whisker stimulation. Right, touch rate during the different contexts. **B**) Firing rate, calculated 0 – 300 ms post first touch, of an example neuron during the different behavioral contexts. **C**) Rasters and histograms of firing rate relative to first touch in example SC and S1 neurons across behavioral contexts. **D**) Mean firing rate in SC and S1 neurons significantly modulated by the Miss condition (133/276 neurons in 6 mice in SC; 75/136 neurons from 4 mice in S1; p < 0.05, Mann-Whitney U-test). **E**) Population histogram of change in firing rate (z-scored) during the Miss condition for all recorded SC and S1 neurons. Neurons that displayed a significant change are denoted in magenta. **F**) Mean firing rate in SC and S1 neurons significantly modulated by the Away condition (102/268 neurons in 5 mice in SC; 30/178 neurons from 3 mice in S1; p < 0.05, Mann-Whitney U-test). **G**) Population histogram of change in firing rate (z-scored) during the Away condition for all recorded SC and S1 neurons. Neurons that displayed a significant change are denoted in green. **H**) Change in stimulus bias in positive preferring SC and S1 neurons during the Away condition (p = 9.5e-12, 268 neurons in 5 mice in SC; p = 9.4e-4, 178 neurons from 3 mice in S1, 2-sided t-test). **I**) Scatter plot comparing the effect of the Away and the Miss conditions in neurons that experienced all three behavioral contexts (158 neurons in 3 mice in SC; 67 neurons from 1 mouse in S1).

### Baseline SC firing rates predict choice probability and speed

Evidence suggests that perceptual decisions are made when neuronal firing rates reach a threshold [26]. Therefore, one potential mechanism for speeding up decision-making is by decreasing the distance of choice-related neurons from their threshold [27]. To examine if “distance to threshold” is modulated by behavioral context, we examined baseline firing rates during Engaged (Hit and CR), Miss, and Away trials (Fig. 6A). On Miss trials, when mice failed to select the upcoming positive stimulus, baseline firing rates in both SC and S1 neurons decreased (Fig. 6B). Interestingly, during Away trials, when the opportunity to select the stimulus was experimentally removed, only SC showed an appreciable decrease in baseline firing rate (Fig. 6C). Differences in locomotion speed across the conditions could not explain these effects (Suppl. Fig. 5A, B). At the single neuron level, SC neurons were more than twice as likely as S1 neurons to display a significant reduction in baseline activity during the Away condition (Fig. 6D). Therefore, baseline SC activity is closely related to choice probability, while S1 activity appears to reflect global changes in task engagement associated with Miss trials. The baseline firing rate of negative preferring SC neurons were largely unaffected by the Away condition, indicating that positive preferring neurons may represent a specialized population of movement related neurons (Suppl. Fig. 5D – F).

**Figure 6.**
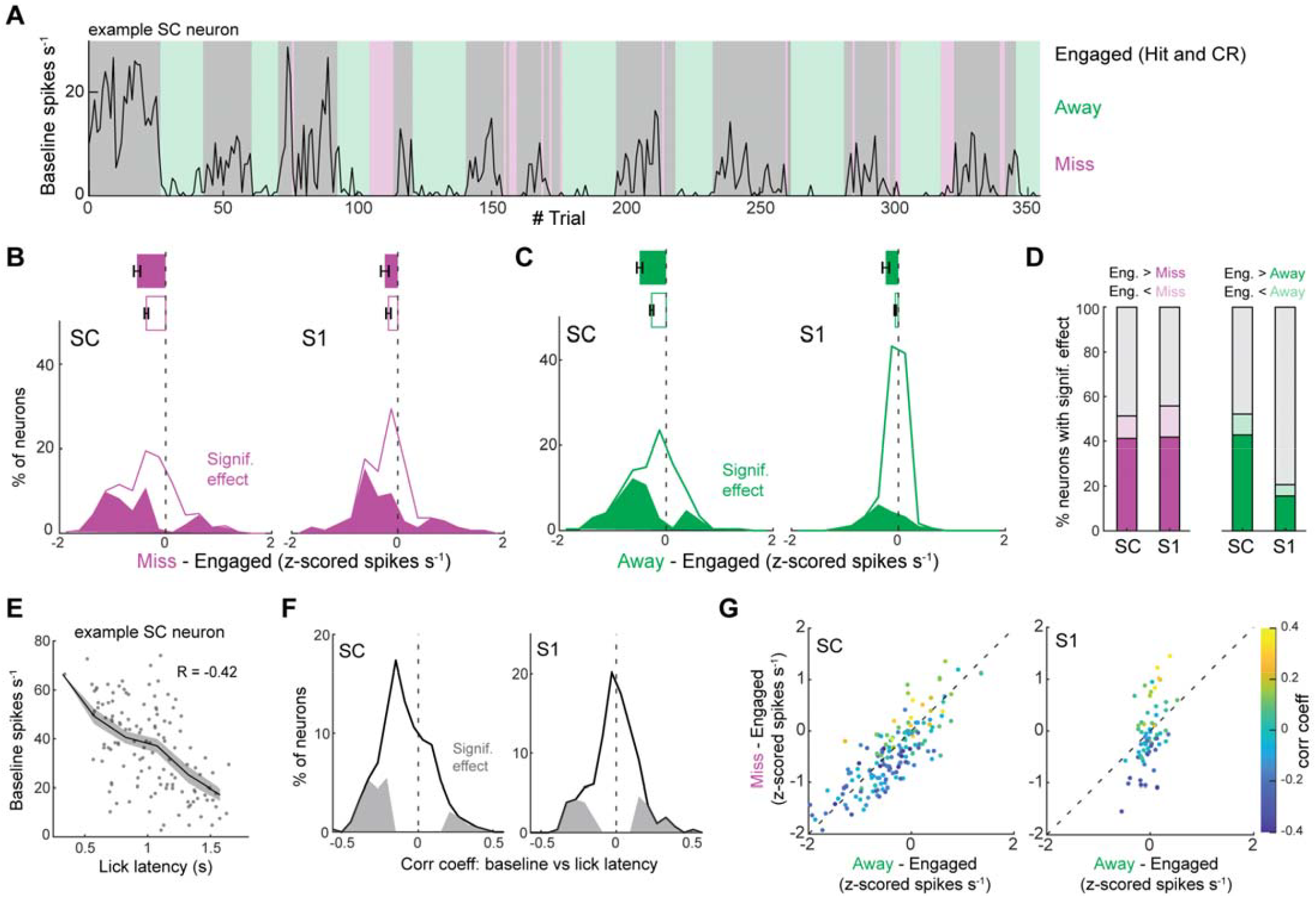
Spontaneous firing rates predict response probability and latency. **A**) Baseline firing rate of an example neuron that during an engaged, away, or miss trial. **B**) Histogram of change in baseline firing rate during the Miss condition. Neurons with a significant effect are denoted in magenta distribution (142/276 neurons in 6 mice in SC; 76/136 neurons from 4 mice in S1; p < 0.05, Mann-Whitney U-test). **C**) Histogram of change in baseline firing rate during the Away condition. Neurons with a significant effect are denoted by the green distribution (140/268 neurons in 5 mice in SC; 37/178 neurons from 3 mice in S1; p < 0.05, Mann-Whitney U-test). **D**) Percentage of neurons with a baseline firing rate significantly modulated by the Miss or Away condition (p < 0.05, Mann-Whitney U-test). **E**) Spearman correlation between baseline firing rate and lick latency in an example SC neuron. **F**) Histogram of the correlation between baseline firing rate and lick latency in the population of SC and S1 neurons (95/386 neurons in 8 mice in SC; 68/263 neurons from 7 mice in S1; p < 0.05, using Matlab *corr* function). Neurons with a significant correlation are denoted by the gray distribution. **G**) Scatter plot comparing the effect of Miss and Away conditions on baseline firing rate. The color axis denotes the correlation of each neuron’s baseline firing rate with lick latency (158 neurons in 3 mice in SC; 67 neurons from 1 mouse in S1).

Next, we reasoned that if SC neurons are involved in driving perceptual choices (i.e. to lick), then their baseline firing rates should correlate to reaction time (Fig. 6E). In line with this reasoning, we saw that higher baseline spike rates in SC but not S1 neurons were associated with shorter response latencies (Fig. 6F). Interestingly, SC neurons with baseline firing rates that were modulated by behavioral context (Miss/Away) were also correlated to response latency, supporting a generalized role in stimulus selection (Fig. 6G).

## Discussion

In this study, we show that primary somatosensory cortex (S1) contains a faithful map of whisker space, where stimulus bias is predominantly controlled by a stable representation of somatotopic space that persists across behavioral contexts. Conversely, we show that superior colliculus (SC) contains a value-based map of whisker space that is strongly biased towards positive stimuli and enhanced by stimulus selection. Moreover, the spontaneous firing rate of positive preferring SC, but not S1, neurons predicted the reaction time of licking. Taken together, these data argue that SC neurons play a key role in transforming somatotopic features into value-based actions. What circuits could be critical to this process?

Both S1 and SC neurons receive input from brain areas involved in sensation, action, and associative learning. In the somatosensory whisker system, the SC gets monosynaptic excitation from trigeminal neurons in the brainstem, while S1 receives brainstem excitation via primary and secondary thalamic nuclei [13,17,28–31]. SC-projecting trigeminal neurons could have greater stimulus bias than their thalamic-projecting neighbors, thereby providing the anatomical substrate for greater stimulus bias in SC. However, whisker-sensitive neurons in the brainstem (SpV) that target the secondary thalamus (Po) also send axons collaterals to SC, indicating a shared ascending sensory representation [32]. If the shared pathway to SC and Po is agnostic to stimulus value, our data suggests that local processing in SC is sufficient for value-modulated somatosensory processing, due to the large bias of the initial (∼20 ms latency) tactile response (Fig. 2B). If the ascending pathway is biased by stimulus value, then a subpopulation of Po→S1 neurons should mirror our results in the SC. This possibility is intriguing, given the known SC→Po pathway that augments tactile responses in S1 [18], and potentially forms a layer-specific value-modulated loop between cortex and midbrain [30,33]. Nonetheless, the contribution of this ascending secondary pathway to sensory maps in S1 remains unclear, yet its activity is known to be modulated by decision-making [34,35]. A technique that enables a cell-type and projection-specific (Po→S1) analysis of S1 neurons is needed to fully resolve this question. Anatomically, the primary pathway from brainstem to S1 is largely distinct from the secondary pathway that involves the SC, yet the primary and secondary brainstem nuclei are known to target both primary and secondary thalamic nuclei, albeit with a clear bias toward their respective targets [1,32]. Ultimately, it remains unclear if the anatomical and functional segregations in the ascending pathways are sufficient to generate the functional differences observed in our study. Future investigations into somatosensory processing in the brainstem are needed to clarify the origin of value-modulated processing in SC and other brain areas.

We discovered that spike suppression on correct rejection trials was a key component of positive stimulus bias in SC neurons. Suppression emerged during the later period (>70 ms) of the sensory response and was preceded by a short window of spike facilitation, suggesting the presence of rapid sensory excitation followed by strong inhibition. SC neurons have large excitatory receptive fields, often facilitated by several different whiskers [36,37]. We hypothesize that the substantia nigra pars reticulata (SNr), a prominent inhibitory input to the SC [38–41], counterbalances this broad sensory excitation to prevent unwanted stimuli from driving orienting movements. This hypothesis is supported by the causal role of the SNr→SC circuit in controlling tongue-mediated target selection [39,42]. Furthermore, the delayed onset of SC suppression in our task mirrors the timing of SNr activation during low-value stimulus presentation [43], and reflects the serial delays imposed by the polysynaptic cortico-striatal pathway [44–46]. Future research focused on dissecting the relative contribution of cortico-striatal and cortico-collicular circuits will provide critical insight into the mechanisms of value-based stimulus selection [39,47–50].

Behavioral context had a significant effect on positive stimulus bias in SC neurons. Stimulus bias significantly decreased when mice were trained to perform detection, where both whisker stimuli were equally associated with reward (Fig. 2). Therefore, the task structure of selecting one particular stimulus amplified bias in SC but not S1. Given this insight, we hypothesized that movement preparation shapes sensory processing in SC neurons. To test this hypothesis, we experimentally removed the opportunity for mice to select the positive stimulus. During this period, mice touched the object similarly but never licked or extended their tongue, indicating they were aware of the experimental constraint. Interestingly, the Hit response in positive preferring SC neurons decreased, leading to a significant decrease in stimulus bias in SC but not S1 (Fig. 5). Therefore, movement preparation increased tactile sensitivity in SC neurons. Similarly, saccade preparation has been shown to increase the sensitivity of monkey SC neurons and enhance task performance in humans [21,51], by presumably directing attention towards the location of the planned movement. In our study, the presence of the water port, which was in close contact with the lips and surrounding hairs, presumably prepared the animal to lick and directed their spatial attention towards the positive whisker. Interestingly, the initial SC sensory response was altered by this movement preparation, indicating that ongoing internal dynamics shape feedforward tactile information in SC neurons. In the visual system, similar effects have been observed not only in SC, but in higher-order visual and parietal cortex [52–54]. It is important to note that movement preparation also modified the later period (100 – 300 ms) of the tactile response, which was unlikely caused by movement initiation, since mice licked with a much longer latency than our analysis window.

A neural correlate of movement preparation was also evident in the pre-stimulus firing rate of SC neurons. When the water port was periodically removed, we observed a significant decrease in baseline spiking exclusively in the positive preferring SC population. With the water port in place, the pre-stimulus firing rate of these neurons was significantly correlated to the animal’s reaction time, suggesting their direct involvement in transforming tactile stimuli into actions. A similar class of SC neurons has been observed in monkeys, where the pre-stimulus firing rate of “build-up” neurons predicted saccadic reaction time [55–57]. This modulation in pre-stimulus firing rates is consistent with the SC mediating a decision threshold [58–61]. At the population level, S1 neurons were largely unaffected by movement preparation. However, during Miss trials, when the animal was presumably disengaged from the task, we also observed a decrease in S1 baseline firing rates. Therefore, task disengagement during Miss trials may coincide with global changes in brain state that spread throughout primary sensory cortex [62–65]. On the other hand, movement preparation may predominantly influence sensory processing in downstream areas more closely associated to action [66].

In this study, we reveal that the representation of sensory space in S1 and SC is differentially modulated by behavioral context. We show that active engagement in value-based stimulus selection greatly biases sensory maps in SC, but not S1, towards the positive stimulus. The circuit mechanisms supporting these differences are unknown, likely involving bottom-up or local changes in feedforward processing as well as top-down influences from cortex and basal ganglia. Understanding the contribution of these circuits to value-based sensory processing will provide critical insight into the neural mechanisms underlying stimulus selection.

## Methods

### Experimental model

This study utilized a total of 23 mice of both sexes with CD1 background. The mice were housed socially in groups of five or fewer per cage. They were kept in a controlled environment with temperatures ranging from 68 to 79°F and humidity levels between 40 and 60%. The mice were maintained on a reverse light-dark cycle (12:12 hours), with experiments conducted during their subjective night. All surgical and experimental procedures were approved by the Purdue Institutional Animal Care and Use Committee (IACUC, 1801001676A004) and the Laboratory Animal Program (LAP).

### Head-plate fixation surgery

Each mouse was fitted with a custom-designed aluminum headplate for head fixation. During the procedure, the animals were anesthetized using 3-5% isoflurane gas. Their body temperature was maintained with a heating pad, and their respiratory rate was continuously monitored to ensure stable anesthesia. To prevent eye dryness during surgery, ointment was applied. The skin and fur overlying the skull were disinfected with 70% ethanol and betadine to minimize the risk of infection. Lidocaine was injected under the scalp and using sterilized surgical instruments, an incision was made along the midline of the scalp. Next, the skin and underlying tissue were carefully removed to expose the skull. To secure the headplate, Liquivet tissue adhesive and Metabond dental cement were applied to the skull and wound edges. Once the headplate was firmly attached and the dental cement had cured, post-operative analgesic was administered, and the mouse was monitored during recovery.

### Behavioral training

Mice were trained in one of two tasks: (i) a whisker discrimination task, and (ii) a whisker detection task. In the whisker discrimination task, one whisker was associated with reward and the other whisker was associated with a punishment. In contrast, in the whisker detection task, both whiskers were associated with a reward of equal value.

Two days after headplate implantation, mice were habituated to running on circular treadmills. This habituation phase consisted of one session per day, lasting 1-2 hours, for 4-7 days. This allowed the mice to become accustomed to locomoting on the circular treadmill. Following habituation, mice were placed on water restriction to increase their motivation for the water reward. Additionally, all whiskers except an adjacent whisker pair (B1 & C1 or C1 & D1) were reduced in length for training on these specific whiskers.

The first stage of training involved classical conditioning. Each trial began with the mouse running two rotations on the circular treadmill, which was followed by the onset of the stimulus. The stimulus consisted of the protrusion of one of two pneumatically controlled touch surfaces (SMC pneumatics), designed to interact with one of the two targeted whiskers (B1 & C1 or C1 & D1) when the mouse was actively sampling the stimulus space. The stimulus remained in the whisker field for 1.5 sec, after which water reward was delivered for the rewarded whisker. After the mice demonstrated anticipatory licking, they progressed to the second stage of training: operant conditioning. In this stage, the water reward was only delivered if the mouse licked in anticipation of the reward following the stimulation of the rewarded whisker. If the mouse licked during presentation of the unrewarded whisker, it was punished by being required to run 2 – 4 times the normal distance to start the next trial.

### Intrinsic signal imaging for localizing whisker barrels in S1

Two days prior to S1 electrophysiology recording, intrinsic signal optical imaging was performed to locate the barrel columns contralateral to the trained whiskers. During the entire imaging session, mice were anesthetized using 1% isoflurane and xylazine (0.3 % mg/kg). The skull over S1 was thinned using a dental drill until the vasculature under it was clearly visible. 100% silicone oil was then applied to headplate well and a cover slip was placed over it to create a flat imaging plane. Under green LED illumination an image of the superficial vasculature was obtained using a CCD camera (Retiga R1™, QIMAGING). Under red LED illumination, changes in fluorescence during whisker stimulus were calculated relative to a baseline period. During the stimulus period, the C1 and D1 whiskers were wiggled using piezoelectric actuators at 20Hz for 4 seconds. Intrinsic signals acquired at a frame rate of 10 Hz over 15 trials with a 10-second intertrial interval were averaged. The sites with greatest change in fluorescence were marked and registered with the superficial vasculature to guide electrode insertion (Supp. Fig 1E). Skull was cleaned and was first covered with a soft silicone gel (Dowsil) and then sealed with a hard-setting silicone (Kwik-Cast™, World Precision Instruments). Mice were monitored for recovery from anesthesia and provided analgesics.

### In-vivo electrophysiology

One day prior to the recording session, mice were briefly anesthetized with isoflurane to perform craniotomies over S1 and SC. For S1, a rectangular window of skull (∼1mm x ∼0.5mm) was carefully removed using a scalpel to expose the brain area overlaying the two whisker barrels columns identified by intrinsic imaging. For SC, a 1 mm circular craniotomy was performed 4 mm posterior and 1.5 mm lateral from bregma. Throughout the procedures, the exposed brain areas were kept moist with ACSF and covered with Dowsil silicone gel and Body Double. On the first day of the recording session, a 128-channel, 3-shank Neuronexus probe was targeted into the S1 barrels identified by intrinsic imaging. The probe was lowered into the cortex at a rate of 75 µm/min using a NewScale micromanipulator. Neural data and experimental signals were acquired at a 20 kHz sampling rate. On the second day of the recording session, the Neuronexus probe targeted the whisker receptive fields in the intermediate and deeper layers of SC. The receptive field of recorded neurons was mapped by deflecting individual whiskers to identify which whiskers elicited the greatest response. If the electrode missed the target, it was repositioned based on somatotopic coordinates.

### Spike sorting

Spike sorting was carried out using the Kilosort2 package in MATLAB, followed by manual curation with the Phy2 GUI (https://github.com/cortex-lab/phy)[67]. To classify spike clusters as single units, we evaluated their spike waveforms, electrode locations, and cross-correlograms to merge, split, or delete clusters as necessary. The spike clusters were validated as signal units based on waveform shapes and auto-correlograms. Only single units were included in all subsequent analyses presented in this paper.

### Trial structure and categorization

The training sessions were conducted in custom-built, sound-attenuated chambers with white background noise and no light. Each session lasted approximately 90 minutes. Trials began after the mouse had locomoted for two rotations on a circular treadmill. Following this, a touch surface protruded into the whisking field of one of the two whiskers, serving as the stimulus. The stimulus was presented for 1.5 seconds. If the mouse licked during this window in response to the rewarded stimulus, a water reward was delivered at the end of the response window. These trials were considered as Hit trials. If the mouse did not lick, it was considered a Miss trial. For a non-rewarded whisker stimulus, if the mouse withheld licking, the trial is considered a correct reject (CR) trial or a false alarm (FA) trial otherwise. To account for satiation and disengagement, non-licking trials which occurred during a consecutive (n>=5) miss trials window are removed from the CR trials.

The Away condition was experimentally controlled via a stepper motor that moved the water spout well outside the reach of the mouse. All experiments started with the spout in the Home position. After 7 positive stimulus trials accumulated in the Home position, the water spout was then moved into the Away position. After 7 positive stimulus trials occurred in the Away position, the spout was returned Home, and the process was repeated. The recording days were the only time the mouse ever experienced the Away condition.

### Whisker tracking & kinematics

The two intact whiskers were imaged at 500 fps for the entire duration of the trial using a high-speed IR camera. DeepLabCut (DLC) was utilized to label the whiskers in each frame, with 4 labels placed on each whisker. To train the DLC network, 200 frames spanning various stimulus conditions for each mouse were manually labelled and used for training the network with over 200k iterations. Whisker angles were calculated for each label with reference to a user-defined point on the face relative to the frame’s vertical axis. Whisker curvature was calculated using the Menger curvature, applied to three points on the whisker at a time, with the mean of all possible combinations taken.

### Identifying touch times

The first onset of touch in each trial was identified as the time the piston entered the whisking field (Suppl. Fig. 1F). To identify individual touch times, a region of interest (ROI) was created at the user-defined interface of the whisker with the object surface. The touch time for each whisk cycle was determined by identifying when the whisker entered the surface ROI. The rare trials where the piston extended behind the whisking field and contacted during the retraction phase of the whisk cycle were excluded from analysis.

### Statistical analyses

#### Stimulus bias

To measure the selectivity of individual neurons to each stimulus, we calculated the stimulus bias at each time point using neuromeric d-prime. It is calculated as the difference in firing rates between two stimuli divided by the root mean square of standard deviation across these stimuli.

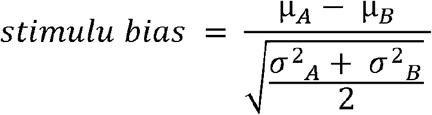

µ_A_ and µ_B_ are the mean spike rates in response to condition A and B respectively. σ^2^_A_ and σ^2^_B_ are the variances of spike rates across trials for condition A and B respectively. This calculation was performed every 10 ms, providing a continuous estimation of bias throughout the response window.

#### Hit and CR preferring neurons

Touch responsive neurons were identified by testing for significant firing rate differences between the baseline period (1.5 sec window pre-touch) and any of the reward/ non-reward stimulus periods (300 ms post-touch) using one-way ANOVA with Tukey post-hoc. To further categorize these neurons based on their stimulus preference, we calculated the mean stimulus bias within the 300 ms window following stimulus onset. Neurons were then determined as either Hit-preferring or CR-preferring based on the direction of this value (positive/negative).

#### Feature Occupancy Control

To control trial-to-trial behavioral variance (run speed and touch strength variance), observations are sub-sampled such that any comparison made across stimulus conditions are from distributions of equal representation of features. Results were randomly subsampled 20 times and averaged from an overlapping feature distribution space.

