## Supplemental figures for "Stimulus selection drives value-modulated somatosensory processing in superior colliculus"

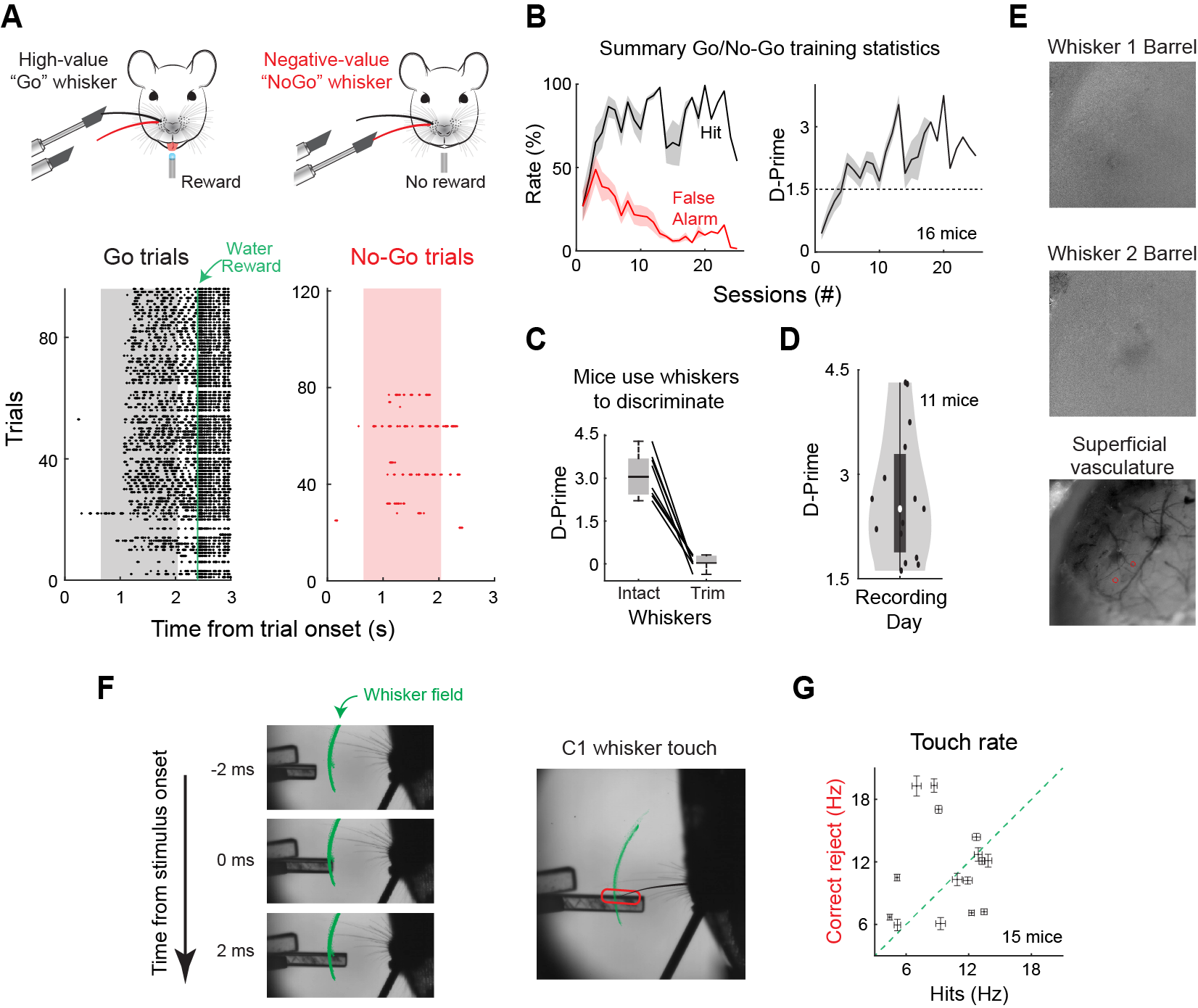

**Supplementary Figure 1. Mice learn value based sensory associations using active whisker touch**

**A**) Example lick rasters from an expert mouse trained to discriminate between C1(Go) and D1(No-Go) whisker stimuli. Shaded region is the stimulus window. **B**) Left: Average hit and false alarm rates over training sessions (n=16 mice). Right: D-Prime over training sessions in the same mice. **C**) D-Prime of expert mice pre and post whisker trim (n = 8 mice). **D**) D-prime of mice during SC and S1 electrophysiology recording (n= 11 mice). **E**) Example images from intrinsic imaging of S1 to locate barrel columns corresponding to whiskers used in the task (Top and middle). The locations of highest intrinsic signal changes overlayed with image of the superficial vasculature to guide electrode placement (Bottom). **F**) Image analysis showing how the onset of first touch was determined from the time the object entered the active whisking field. **G**) Mean touch rate for first 300ms of sensory period during hit and correct reject stimuli (n = 15 mice).

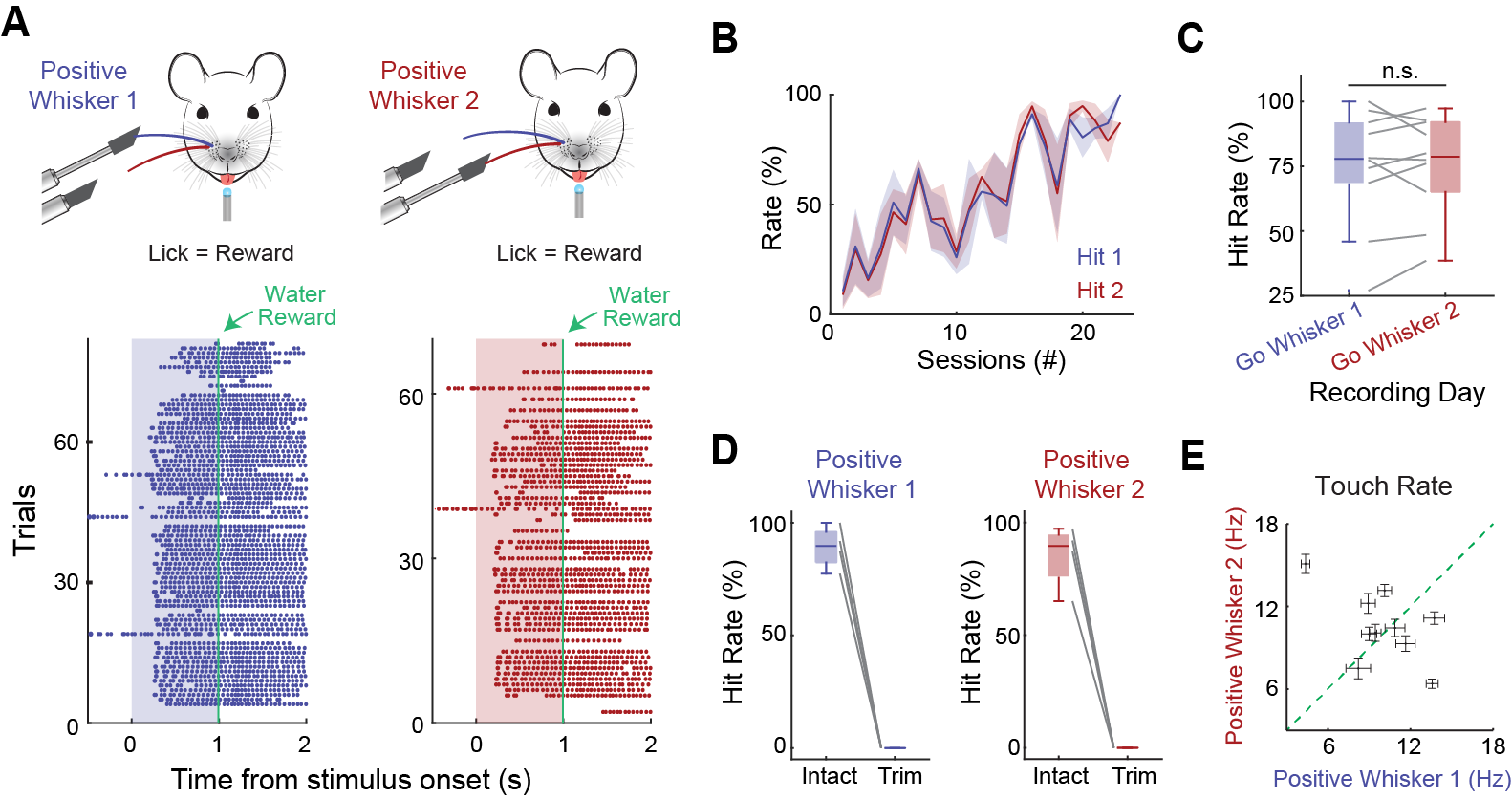

**Supplementary Figure 2. Mice learn equal valued sensory associations.**

**A**) Example lick rasters from an expert mouse trained to lick in response to either of the two single whisker stimuli.

**B**) Average hit rates for both single whisker stimuli over training sessions (n = 7 mice). **C**) Hit Rates during S1 and SC electrophysiology recordings (n = 6 mice). **D**) Hit rates during both single whisker stimuli pre and post whisker trimming (n = 4 mice). **E**) Mean touch rate for first 300ms of sensory period during positive whisker 1 and 2 stimuli (n = 10 mice).

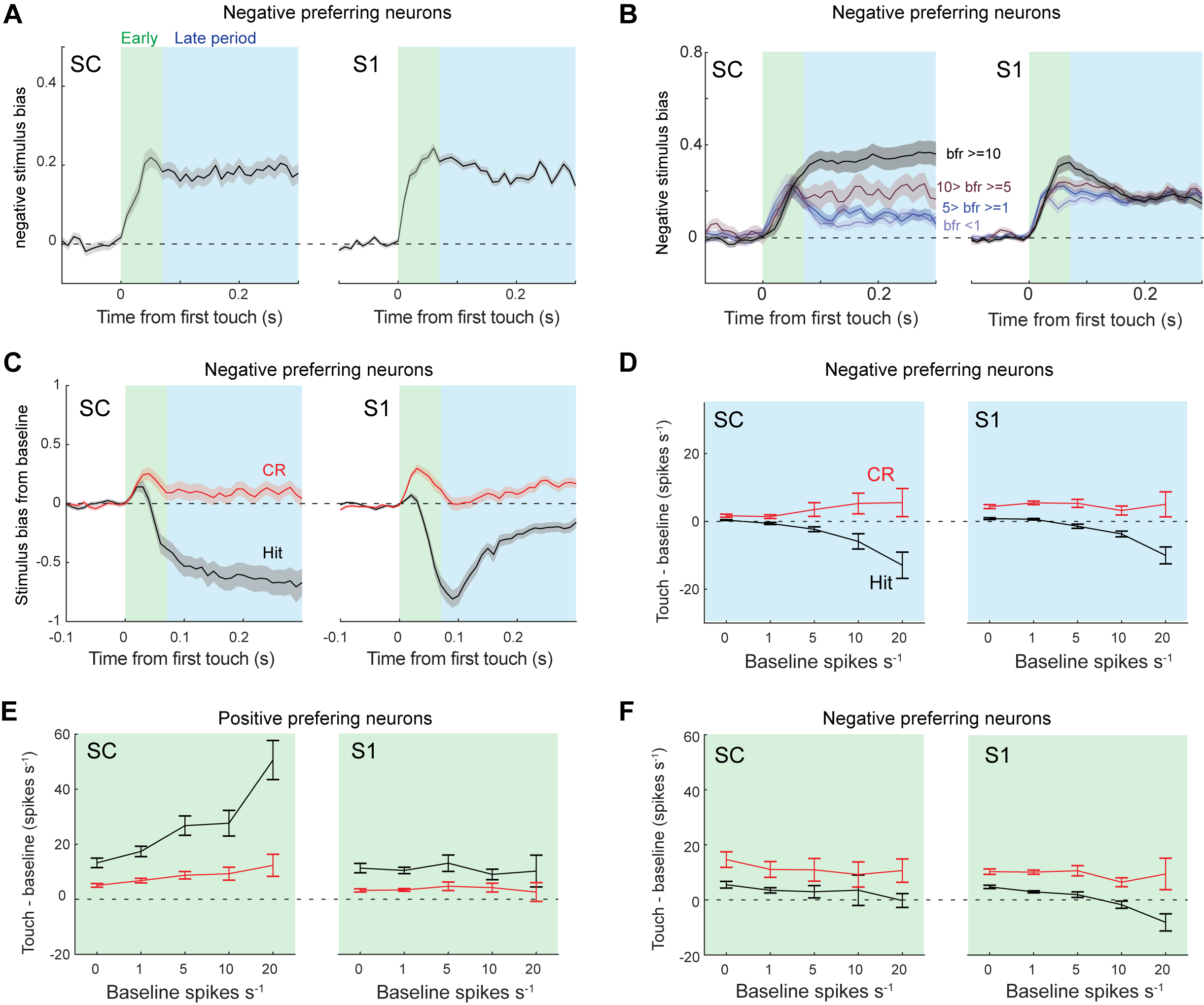

**Supplementary Figure 3. Stimulus of negative preferring neurons are similar between SC and S1.**

**A**) Negative stimulus bias in neurons that preferred the negative valued stimulus. (163 neurons in 8 mice in SC; 362 neurons from 7 mice in S1). **B**) Same as in A except with the data grouped by baseline firing rate. (For SC: 51 neurons in 8 mice in bfr>=10 group; 24 neurons in 7 mice in 10>bfr>=5 group; 48 neurons in 8 mice in 5>bfr>=1 group; 40 neurons in 8 mice in bfr<1 group; For S1: 78 neurons in 7 mice in bfr>=10 group; 64 neurons in 7 mice in 10>bfr>=5 group; 149 neurons in 7 mice in 5>bfr>=1 group; 71 neurons in 7 mice in bfr<1 group). **C**) Bias from baseline in negative preferring neurons. (79 neurons in 8 mice in SC; 164 neurons from 7 mice in S1; Showing neurons with baseline spike rate > 4). **D**) Mean touch-evoked firing rates during the late period of Hit and CR response in negative preferring neurons. **E**) Mean touch-evoked firing rates during the early period of the Hit and CR response in positive preferring neurons. **F**) Mean touch-evoked firing rates during the early period of the Hit and CR response in negative preferring neurons.

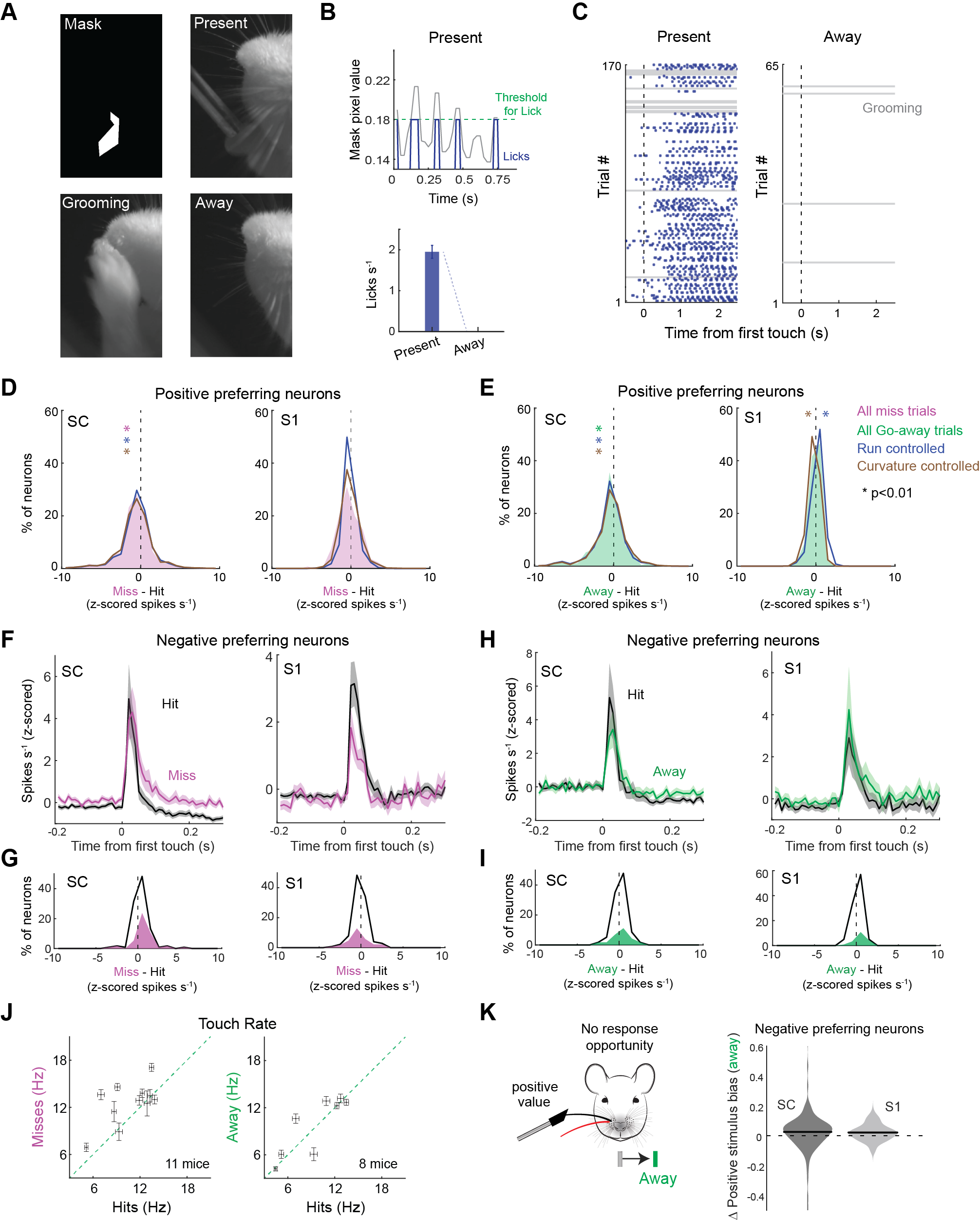

**Supplementary Figure 4. Unexpected licking, locomotion, or touch quality cannot explain the effects of the Miss or Away conditions. A**) Image frames of facial movement analysis for determining if the animal licked when the water port was Away. **B**) Quantification of image pixel values for extracting lick rate during present and Away conditions. **C**) Lick and grooming raster for the Present and Away conditions. **D**) Change in spike rate in positive preferring neurons for all trials and for trials where the locomotion speed and curvature were subsampled to match the occupancy of the Miss condition (For SC: All trials p = 2.60e-8; Run controlled: p = 8.53e-6; Curvature controlled: p = 5.51e-7; For S1: All 3 sampling conditions have p > 0.01, unpaired t-test). **E**) Same as D, except for the Away condition (For SC: All trials p = 6.51e-7; Run controlled: p = 8.22e-7; Curvature controlled: p = 4.02e-4; For S1: All trials p >0.01; Run controlled: p = 1.33e-3; Curvature controlled: p = 1.40e-6, unpaired t-test). **F**) Firing rates of negative preferring neurons during Hit and Miss trials (53/163 neurons in 6 mice in SC; 84/362 neurons from 4 mice in S1; p < 0.05, *Mann-Whitney U-test*). **G**) Histogram of change in neuronal spike rates during the Miss condition. **H**) Same as F, except for the Away condition (32/117 neurons in 6 mice in SC; 23/121 neurons from 4 mice in S1; p < 0.05, *Mann-Whitney U-test*). **I**) Same as G, except for the Away condition. **J**) Scatter plot of mean ± SEM touch rates calculated during Hit, Miss, and Away trials of each mouse in the study. **K**) Change in positive stimulus bias in negative preferring neurons during the Away condition (p = 0.0401, 117 neurons in 5 mice in SC; p = 7.06e-5, 121 neurons from 3 mice in S1*, 2-sided t-test*).

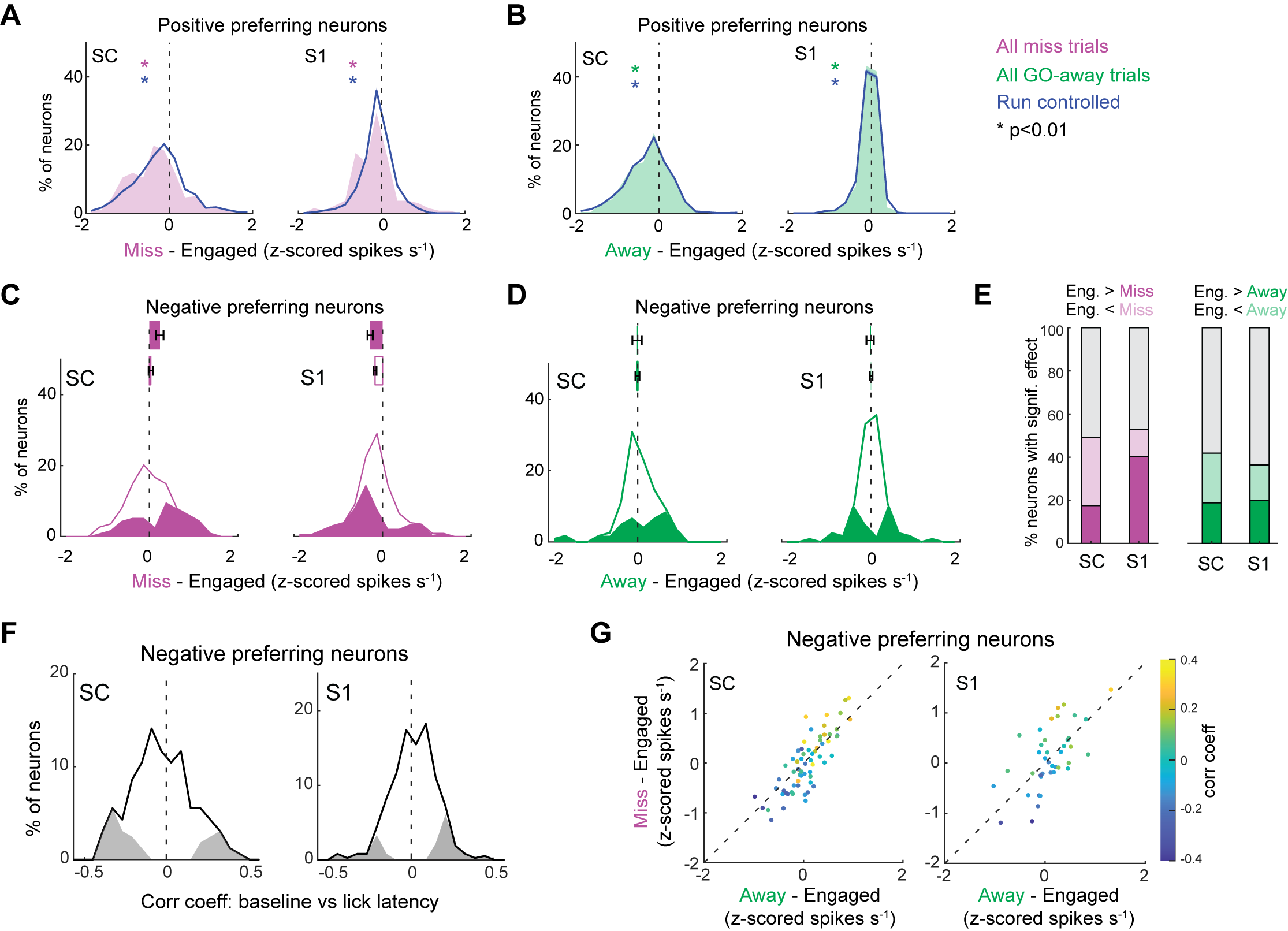

**Supplementary Figure 5. Baseline firing rate: locomotion speed control and the effect of behavioral context on negative preferring neurons. A**) Change in baseline firing rate in positive preferring neurons, comparing the Miss effect calculated from all trials to the locomotion occupancy-controlled Miss effect (For SC: All trials p = 1.01e-20; Run controlled: p = 1.89e-14; For S1: All trials p = 1.24e-4; Run controlled: p = 3.72e-4, unpaired t-test). **B**) Same as in A, except for the Away condition (For SC: All trials p = 6.90e-15; Run controlled: p = 7.64e-16; For S1: All trials p = 7.95e-4; Run controlled: p = 5.87e-4, unpaired t-test). **C**) Histogram of the change in baseline firing rate in negative preferring neurons during the Miss condition (56/114 neurons in 6 mice in SC; 122/231 neurons from 4 mice in S1*;* p < 0.05, *Mann-Whitney U-test*). **D**) Same as in C, except for the Away condition (49/117 neurons in 5 mice in SC; 44/121 neurons from 3 mice in S1*;* p < 0.05, *Mann-Whitney U-test*). **E**) Bar graph of the percent of neurons that significantly increase, decrease, or do not change during the Miss or Away condition (p < 0.05, *Mann-Whitney U-test*). **F**) Correlation coefficient between baseline spike rate and lick latency in negative preferring neurons (41/163 neurons in 8 mice in SC; 65/362 neurons from 7 mice in S1*; p < 0.05, using Matlab* corr *function*). **G**) 3-dimensional scatter plot comparing the effect of Miss and Away on baseline spike rate and the correlation coefficient with reaction time in negative preferring neurons (*68* neurons in *3* mice in SC; *40* neurons from *1* m*ouse* in S1).
